# Spatial and temporal representation of marine fish occurrences available online

**DOI:** 10.1101/2023.03.09.531981

**Authors:** Vanessa Pizarro, Andrea Castillo, Andrea Piñones, Horacio Samaniego

## Abstract

Despite the 243,000 species of marine species described by 2022, our knowledge about the biodiversity in the oceans is still incomplete. This may have dreaded and detrimental effect for the conservation of marine ecosystems under the current anthropization of our biota and the fast pacing climate and global change scenario.

However, a large number of online repositories cataloging, storing and distributing biodiversity information hosting taxonomic information and species occurrence data have emerged recently. FishBase, the Global Biodiversity Information Facility (GBIF) and the Ocean Biodiversity Information System (OBIS) are part of these publicly available repositories representing a variety of sources that have exploded in number. However, despite the incredible accumulation of biodiversity records, not all the information is really useful, nor does it represent any new knowledge regarding global species richness patterns.

In this study we evaluated the spatial and temporal representativeness of the records of marine fishes (order Actinopterygii) available in the GBIF and OBIS global repositories. We provide a methodological framework based on a set of non-parametric estimators to calculate species richness from incidence data, using hexagonal grids as sampling units overlapped on the marine bioregions worldwide.

Using standard ecological and spatial analysis tools, we identify regions that are adequately represented in terms of available records and therefore have more reliable data, as well as regions with few records that do not represent current species richness. We overlap these results with the location of marine protected areas and fishing exploitation zones to understand the anthropogenic effect on marine ichthyofauna. We additionally evaluate hypotheses regarding the taxonomic, geographic, and temporal distribution of information biases to deepen our current understanding of public records of species occurrences worldwide.

Considering that more than 40 years of information was analyzed, the results showed that on a global scale, the primary data on marine fish available on GBIF and OBIS platforms are still far from being representative and complete. Only 1.14% of the records were useful for our analyses. In addition, we found that the information seems to be biased towards coastal areas, regions close to developed countries and areas where there is a large fishing activity. Finally, the best represented species and families are those with a small body size, which use shallow habitats and have commercial cultural value.

## 1. Introduction

Currently, the more than 243,000 species included in the World Register of Marine Species database (WORMS, 2022) suggests that only 11 to 78% of all marine species have been discovered revealing a striking picture of vastly incomplete knowledge that may have serious implications for marine conservation (Luypaert et al., 2020). Moreover, ongoing climate change represents one of the greatest threats to biodiversity (Malhi et al., 2020; Turner et al., 2020) and has already been documented to modify the distribution of marine species (Lenoir et al., 2020). Some of the described effects considers the invasion of non-native species leading to massive species turnover that may lead to the local extinction of large proportions of species (Cheung et al., 2009).

While species richness is often used to represents diversity patterns, species richness is, in itself, an aggregate variable subsuming the variety of life (Marquet et al., 2004). Hence, several attempts have focused on the development of more encompassing indices sparking interesting scientific debates to describe ecological heterogeneity (Tuomisto, 2011; Moreno and Rodríguez, 2011; Daly et al., 2018). However, scientific literature seem to have opted to shift focus to the consequences of biodiversity loss by fostering the usage of new terminology to provide hands-on concepts designed to convey messages to policy makers summarizing the richness of biodiversity (Pereira et al., 2013; Butchart et al., 2010). Still, while scientists have debated the use biodiversity terminology, species richness provides a succinct, and easy-to-handle description of the variability across several other quantities describing biodiversity in space and time (Appeltans et al., 2012), and is, together with other diversity indices, an essential feature to understand how diversity changes under the impact of natural and anthropogenic factors on biomes, regions and ecosystems (Troia and McManamay, 2017; Magurran and McGill, 2011).

Likewise, biodiversity can also be assessed through life history traits, which are modulated by both evolutionary factors and the variation in habitats and ecosystems (Neigel, 1997; Hutchings and Baum, 2005). We now know that biodiversity is more likely an expression of the heterogeneity of such life history traits. Alo et al. 2021, for example, shows that while some of the fish diversity is certainly due to environmental processes, a large fraction of such richness variance is also determined by evolved life history traits related, for example, to migratory habits. Therefore, evaluating how life history traits impact richness metrics should deepen our understanding of fish diversity patterns.

While still short of having a robust and standardized biodiversity infrastructure (Heberling et al., 2021), a great diversity of online repositories with taxonomic information and species occurrences data exist. Among the most important databases hosting marine information are FishBase, a platform that hosts information on the taxonomy of fish, their ecology, trophic information, habitat and history of uses dating back to more than 250 years (Froese and Pauly, 2000); the Global Biodiversity Information Facility (GBIF), a platform that stores and allows for the free access to species occurrence records from around the world. GBIF is currently one of the repositories hosting the largest amount of such data in the world (Telenius, 2011; GBIF: The Global Biodiversity Information Facility, 2021); and finally Ocean Biodiversity Information System (OBIS), which houses data on the occurrence and abundance of species from exclusively marine environments (OBIS: Ocean Biodiversity Information System, 2021). Records entered in these repositories are often used for research related to biodiversity assessment, taxonomic reviews, red listing of threatened species, species distribution, and generation of ecological niche models, among others (Yesson et al., 2007). GBIF currently offers more than 1.62 billion occurrence records and OBIS more than 63 million, which increase considerably each year (GBIF: The Global Biodiversity Information Facility, 2021; OBIS: Ocean Biodiversity Information System, 2021).

The records of both platforms come from a wide variety of sources collected following different methodologies at different temporal and spatial scales introducing a great variety biases (Beck et al., 2014; Zizka et al., 2020). Amon these, three main types of biases have been described: (i) taxonomic, this occurs when some species and/or families are better sampled than other rarer species (Chandler et al., 2017); (ii) geographic, when data input is unevenly distributed across geographic regions and may prove to obscure inter-region comparisons (Yang et al., 2013; Yesson et al., 2007); and (iii) temporal, which may be prevalent when comparing different time periods as data coverage is unevenly distributed over time (Chandler et al., 2017; Yang et al., 2013). While these biases introduce some uncertainty regarding reliability of species richness descriptions obtained from online platforms (Beck et al., 2014; García-Roselló et al., 2015), they have largely been used to provide an extensive overview of macro-ecological patterns of distribution not available otherwise (Mora et al., 2008; Troia and McManamay, 2017).

Still, identifying how sampling effort is distributed across space and time will help to interpret biodiversity patterns and reduce biases. This may be achieved through different weighting schemes for records in areas with sufficient sampling that provide a more reliable contribution compared to underrepresented regions (Phillips et al., 2009; Hortal et al., 2008; Yang et al., 2013).

We here assessed the spatial and temporal representativeness of marine fish records available in the global GBIF and OBIS repositories at the level of marine bioregions in order to pinpoint the location of records that best quantify the diversity of marine fishes. The result is a spatial representativeness analysis that we then overlay on marine conservation areas (UNEP-WCMC and IUCN, 2022) and fisheries exploitation areas (FAO, 2014) to learn whether marine conservation efforts, as well as large fisheries, are located in areas of high species richness or in areas of insufficient data coverage.

Finally, we also analyzed the potential effect that some attributes could have on the incidence of more records in global database repositories. Specifically, we evaluated three hypotheses related to body size, habitat depth and commercial use. The underlying hypotheses are that a better representation in the online platforms may be due to the over sampling of larger fish, caused by its easy identification; that shallow areas provide easy access to sampling; and economic and commercial interest have elicit a larger representation of culturally relevant species among online biodiversity repositories.

## 2. Methods

### 2.1. Species data

We use all recorded ocurrences from the order Actinopterygii hosted in GBIF and OBIS repositories (GBIF.org, 2021; OBIS.org, 2021). Following Alò et al. (2021), evolutionarily older taxa, such as Cephalaspidomorphi, were excluded from this analysis. The libraries *rgbif* and *robis* of the statistical package R were used for data extraction (Chamberlain, 2017; Provoost and Bosch, 2020; R Core Team, 2018). To minimize errors associated with the public usage of GBIF and OBIS repositories, we curated the dataset following Zizka et al. (2020) and filtered the dataset by the columns labeled “scientific name”, “family”, “year”, “Longitude” and “Latitude”. We retained all taxonomic information down to the species level. Any record with NA values was removed. We also removed any duplicated record with identical latitude and longitude as well as any record collected before 1980 (see Alò et al., 2021; García-Roselló et al., 2015). Each record was further assigned to a marine bioregions following Costello and Chaudhary (2017). Spatial data wrangling and plotting was performed with the aid of the following libraries: *sf, dplyr*, and *cartography* (Giraud and Lambert, 2016; Pebesma, 2018; Wickham et al., 2021). We finally labeled and removed any exotic species record using the distribution() function provided by the *rfishbase* library (Boettiger et al., 2012; Froese and Pauly, 2021). To limit our analysis to species occurring within their native range each record was checked against the classification of FAO fisheries area for consistency (FAO, 2014).

### 2.2. Data Analysis by Bioregion

Once the database was cleaned, a subset of the data was created for each of the 30 bioregions. For each bioregion, a count of records, species and families was made, and the Shannon diversity index was calculated using the *vegan* library in R (Oksanen et al., 2020).

### 2.2.1. Spatial Representativeness Analysis

To assess the spatial representativeness of the data, bioregions were gridded into hexagonal cells of equal area to maximize fit to bioregions areas. These 1°× 1° hexagonal lattice yielded 60,853 cells in total. We evaluated, in the appendix, two additional spatial resolutions, of 5° and 10° lattice with a total of 3,021 and 958 cells respectively, in order to analyze different biodiversity macropatterns (Tittensor et al., 2010). The expected species richness (*S*_*exp*_) was computed as the average between three non-parametric richness estimators so that 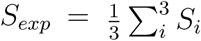, where *S*_*i*_ is Chao2 (*S*_*chao*_), Bootstrap (*S*_*bootstrap*_) and Jackknife 1 (*S*_*jackknife*1_) (see Magurran and McGill, 2011, for individual definition of indices). Such averaging seeks to minimize biases and potential errors of under- or over-estimation by using a single richness estimator following the work of (Mora et al., 2008; Troia and McManamay, 2017).

We then produced a spatial representativeness index (SRI) by comparing the observed richness (*S*_*obs*_) per cell to *S*_*exp*_ (Troia and McManamay, 2017).

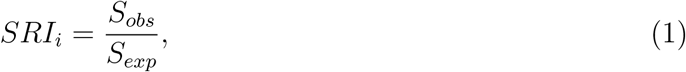

indicates the degree of representativeness of records to quantify the actual species richness in each cell (*i*). Its value ranges from 0 to 1, where 0 represents an unsampled cell and 1 represents a fully sampled one (Eq. 1).

Because SRI somehow shows how databases depict the actual species richness, we may further classify SRI into four classes labeled by the levels of species richness knowledge they represent. Some cell may show very *few* knowledge with *SRI ∈* (0, 0.60). Conversely, others may show to have a *sufficient* species diversity knowledge level for a complete representation of species diversity if *SRI ∈* (0.60, 0.85). While others, in turn, will have an *adequate* representativeness level if *SRI ∈* (0.85, 1.00). Cells with *no records* were also considered as an independent class.

### 2.2.2. Temporal Representativeness Analysis

We generated plots of cumulative records over time to analyze the temporal distribution of data records for each bioregions. Accumulation curves for the 30 bioregions were calculated based on the observation records and the year of collection. We asses the completeness of the sample by evaluating the final 10% tip of the curve using a linear fit after rescaling *S* to create statistically comparable slope units. Slopes close to 0 indicate sufficiently sampled bioregions, while slopes closer to one are indicative of insufficient sampling efforts across time.

### 2.2.3. Gap Analysis

We overlaid the spatial representativeness map (§2.2.1) with shapefiles of Marine Protected Areas (MPA) (UNEP-WCMC and IUCN, 2022) and fishing exploitation areas reported by (FAO, 2014). The superposition of these layers allowed us to calculate the extent of protection offered by MPA for each bioregions on a cell basis, and the extent of cells in designated fishing zones. This exercise allows to jointly assess the relationship between two opposing human impacts and current uncertainties about marine fish diversity.

### 2.2.4. Bias Assessment

The evaluation of potential biases generated by body size, habitat depth and cultural value of species (§2.1) was assessed from the fishbase database (Froese and Pauly, 2021). Size frequencies were determined using 80 cm intervals and ranges of habitat depth were determined according to the classification of oceanic layers (i.e. epipelagic = 0 - 200 m, mesopelagic = 200 - 1,000 m, and bathypelagic= 1,000 - 4,000) (Costello et al., 2010). Parametric correlation analysis was employed describe the relationship between the frequency of representation, using a logarithmic transformation, and the body size and habitat depth, while a a simple pie chart shows the frequency of cultural values associated to the data.

All data and scripts are available (see Appendix A).

## 3. Results

### 3.1. Records by Bioregions

Just about 1.14% of the total published occurrences in the order Actinopterygii were retained in our analysis. That is, from the 71,670,596 downloaded records off the GBIF and OBIS repositories, 820,004 were considered useful (see Appendix A). This subset consisted of 10,371 species in 361 families. The most represented families in our dataset are Scombridae, Pleuronectidae and Gadidae with 103,762, 57,018 and 52,079 records respectively. The species with the largest representation frequency are *Hippoglossoides platessoides, Mola mola* and *Coryphaena hippurus* with 30,885, 21,042 and 21,089 records.

The analysis at bioregion level (Table 1) shows a large variability. The count of records varies across three orders of magnitudes, that is from 2.68 × 10^5^ records in the Caribbean Sea and Gulf of Mexico (11) down to 1.02 × 10^2^ in the Black Sea (2). The bioregion with the largest species richness and diversity index is the the Indo-Pacific Seas and Indian Ocean (13) with 2.95 × 10^3^ recorded species and a Shannon index of 6.93 followed by the Coral Sea bioregion (16) with 2.93 × 10^3^ species and a Shannon index of 6.75. Likewise, the Coral Sea also presents the largest number of families. It is interesting to note that, while being the largest bioregion (i.e. in km^2^), the Southern Ocean show the fewest number of records and the lowest number of species and families across all bioregions. Black Sea (2) and Norwegian (4) are the bioregions with lowest number of record and Shannon index value respectively. Fig. 1 illustrates the location of the 30 marine bioregions and their respective richness and diversity values.

**Table 1:** Counts of records, species, families and Shannon diversity for each bioregion. The largest values are highlighted in bold.

| ID | Bioregion | Area (km<sup>2</sup>) | Records | Species | Families | Shannon |
| --- | --- | --- | --- | --- | --- | --- |
| 1 | Inner Baltic Sea | 415,265 | 8,902 | 72 | 30 | 2.46 |
| 2 | Black Sea | 537,390 | 102 | 37 | 22 | 3.21 |
| 3 | NE Atlantic | 2,053,367 | 87,377 | 310 | 104 | 3.90 |
| 4 | Norwegian Sea | 1,132,035 | 3,046 | 93 | 35 | 2.16 |
| 5 | Mediterranean | 2,859,080 | 12,532 | 372 | 101 | 3.39 |
| 6 | Arctic Seas | 10,276,384 | 2,506 | 114 | 23 | 3.90 |
| 7 | North Pacific | 12,974,899 | 78,070 | 839 | 156 | 4.50 |
| 8 | North American boreal | 8,001,411 | 9,709 | 162 | 48 | 2.99 |
| 9 | Mid-tropical N Pacific Ocean | 32,685,992 | 9,310 | 615 | 127 | 4.59 |
| 10 | South-east Pacific | 21,952,036 | 386 | 190 | 89 | 4.97 |
| 11 | Caribbean and Gulf of Mexico | 8,427,282 | **268,066** | 1,703 | 209 | 4.49 |
| 12 | Gulf of California | 6,184,523 | 7,639 | 885 | 148 | 5.93 |
| 13 | Indo-Pacific seas and Indian Ocean | 37,090,022 | 16,967 | **2,947** | 215 | **6.93** |
| 14 | Gulfs of Aqaba, Aden, Suez, Red Sea | 830,183 | 926 | 352 | 72 | 5.51 |
| 15 | Tasman Sea | 3,592,495 | 1,003 | 380 | 120 | 5.36 |
| 16 | Coral Sea | 7,658,020 | 40,107 | 2,929 | **249** | 6.75 |
| 17 | Mid South Tropical Pacific | 23,418,352 | 6,083 | 811 | 123 | 5.18 |
| 18 | Offshore and NW North Atlantic | 16,012,294 | 130,994 | 897 | 190 | 3.46 |
| 19 | Offshore Indian Ocean | 31,076,432 | 1,263 | 337 | 116 | 4.06 |
| 20 | Offshore W Pacific | 10,291,385 | 6,363 | 1,839 | 232 | 6.81 |
| 21 | Offshore S Atlantic | 41,435,619 | 11,960 | 990 | 188 | 3.79 |
| 22 | Offshore mid-E Pacific | 13,815,068 | 687 | 79 | 37 | 3.04 |
| 23 | Gulf of Guinea | 3,325,856 | 6,816 | 384 | 138 | 3.95 |
| 24 | Argentina | 2,665,312 | 8,701 | 115 | 52 | 2.83 |
| 25 | Chile | 1,739,182 | 250 | 100 | 54 | 4.36 |
| 26 | Southern Australia | 3,824,338 | 15,643 | 1,011 | 201 | 5.75 |
| 27 | Southern Africa | 4,371,199 | 19,954 | 1,142 | 210 | 4.16 |
| 28 | New Zealand | 6,293,426 | 53,879 | 558 | 154 | 3.66 |
| 29 | North West Pacific | 2,457,257 | 1,767 | 869 | 182 | 6.46 |
| 30 | Southern Ocean | **62,161,808** | 8,996 | 294 | 57 | 3.98 |

**Fig. 1:**
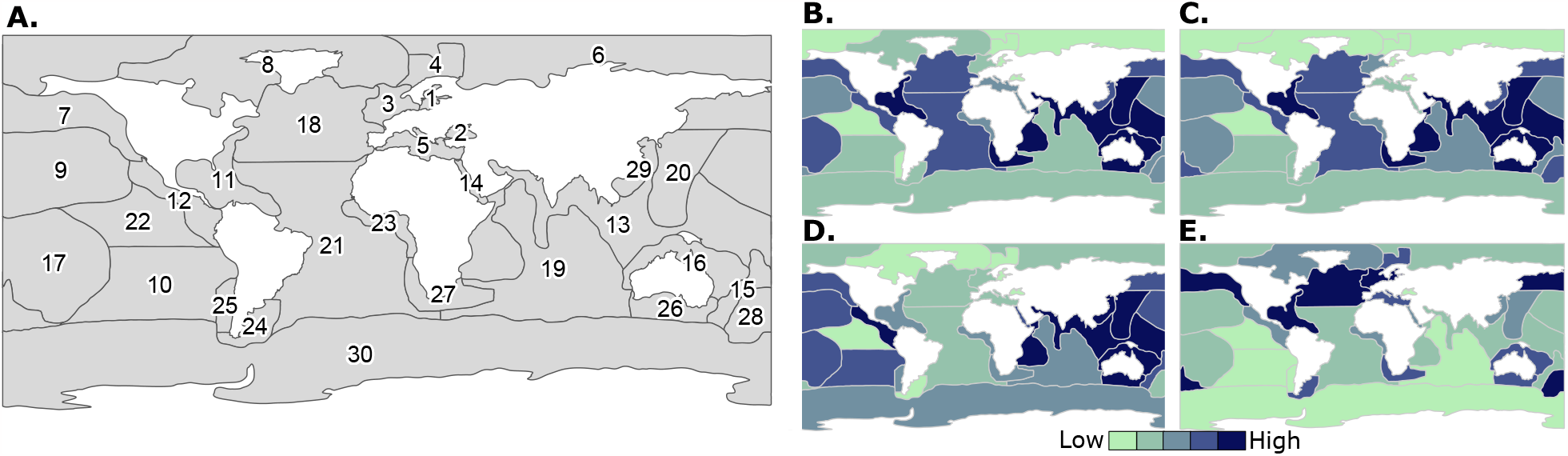
Marine bioregions and spatial diversity distribution used in this study. **A**. The 30 marine bioregions from Costello and Chaudhary (2017) used in this study. Number are identification labels in Table 1. **B**. Overall species richness across bioregions; **C**. Family richness; **D**. Average richness (see §2.2.1); and **E**. Shannon diversity index. Note that values in **B-E** have been normalized for display purposes. See Table 1 for actual values.

### 3.2. Geographic Analysis

Fig. 2 shows the cell classification according to SRI (§2.2.1). As expected, no bioregion is completely sampled at the 1° scale resolution. In fact, at this resolution scales, large empty regions with no records are observed. The bioregions with the largest area classified as *Adequate* are the Northeast Atlantic (3) (38.6%), the Caribbean and Gulf of Mexico (11) (22.8%) and the Inland Baltic Sea (1) (21.8%). It should be noted that such cells mainly correspond to coastal areas in the northern hemisphere. On the other hand, the bioregions that present a greater surface without records correspond to the Southeast Pacific (10) (97.5%), the Southern Ocean (30) (95.7%) and the South Pacific, Mid Tropical (17) (93.9%). Additional results for 5°× 5° and 10° × 10° spatial resolution grids are available in Appendix B.

**Fig. 2:**
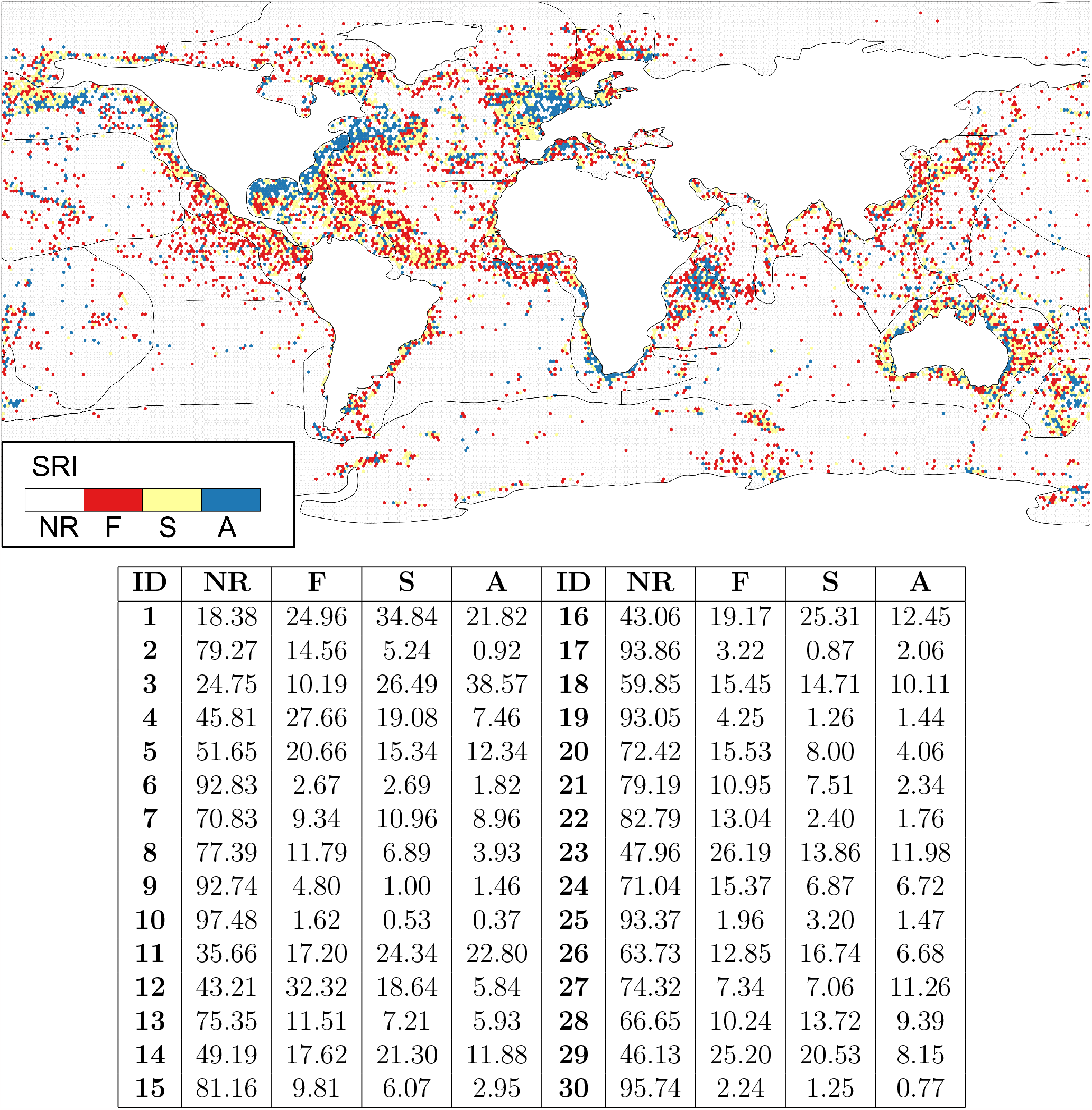
Spatial Representativeness Index (SRI) in 1° hexagonal lattice. Values in the table below indicates the surface area as a percentage of each bioregion for every SRI category (see §2.2.1). **ID** is the identification number given to each bioregion (Table 1). **A** are cells with an adequate representativeness of species richness (i.e. SRI *>* 0.85. **S** are cells considered as having a sufficient representativeness (i.e. SRI *∈* (0.60, 0.85). **F** cells are cells with few records and are thus not considered to be representative of actual species richness (i.e. SRI *∈* (0, 0.6)). **NR** as cells with no records (SRI= *NA*).

### 3.3. Temporal Analysis

Bioregions show similar trends of data accumulation across the four decades analyzed here (Fig. 3). While a significant increase is apparent in the time period between 2005 and 2010, such increase is not significant for 14 out the 30 bioregions. The Caribbean and Gulf of Mexico

**Fig. 3:**
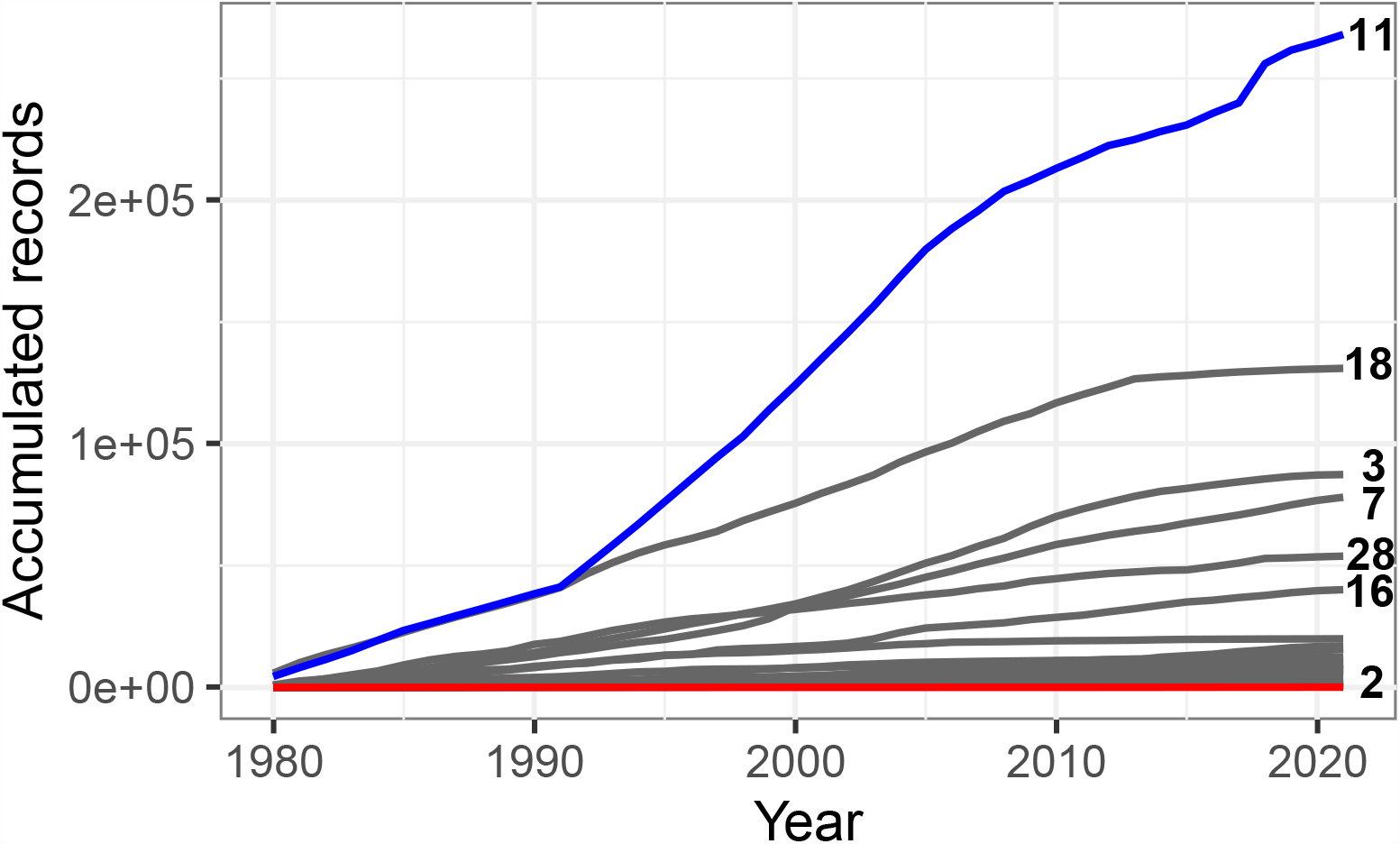
Records accumulation rate for each bioregion across the four decades analyzed. The blue line is the accumulation of fish records in the Caribbean and Gulf of Mexico bioregion (11) and the red line shows the accumulation rate in the Black Sea (2). Numbers as the end of each timeseries correspond to the bioregion ID in Table 1.

(11) is the bioregion with the largest increases in data contribution to the dataset, while the Black Sea (2) is the bioregion with the lowest rate of data contribution in the 40 years span between 1980 and 2020. (See Appendix C for further analysis).

We categorize the slopes of the final 10% of each accumulation curves in Fig. 4. Fourteen bioregions show a slope less than 1. The Mediterranean Sea (5) stands out with the lowest slope value (0.47), while the Black Sea (2) is the bioregion with the steepest final slope (3.13).

**Fig. 4:**
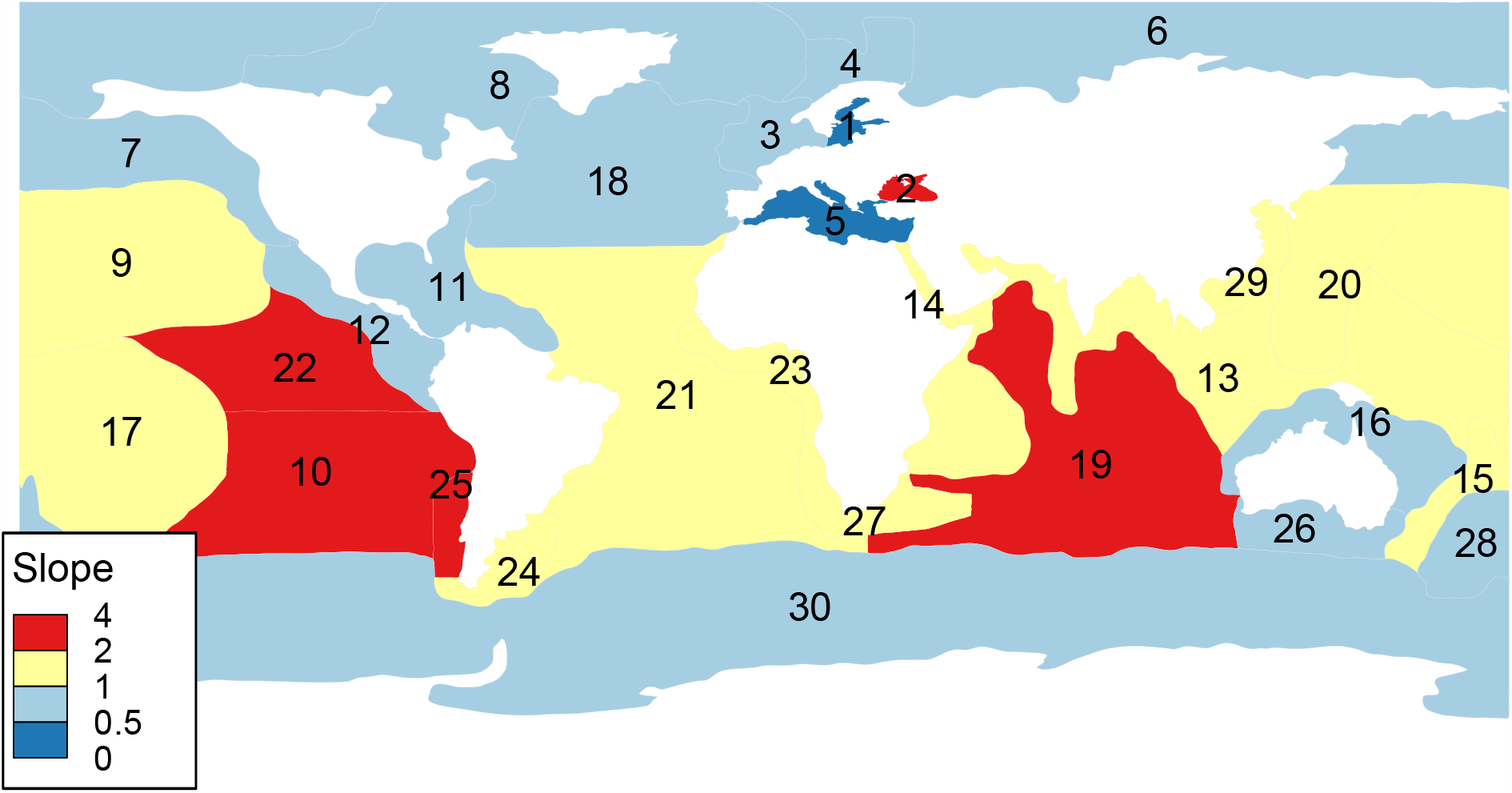
Graphical representation of the slope values of the species accumulation curve for each bioregion. The slope corresponds to the final 10% of the species accumulation curve. See §2.2.2 for details regarding the analysis.

### 3.4. Gap analysis and fishing exploitation areas

Most PAs are concentrated in the North Atlantic bioregions close to the European continent (Table 2). The largest incidences of PAs across bioregions are for the Northeast Atlantic (3) followed by the Inner Baltic Sea (1) (n=296) and the Mediterranean Sea (5) (n=214) with 734, 296 and 214 PAs respectively. However, the bioregions with the largest area covered by protected areas are the Coral Sea (16), the northeast Atlantic (3) and New Zealand (28) covering a 37.3, 17.4 and 16% of their respective areas. Regarding the sampling level of these bioregions, the Northeast Atlantic (3), the offshore and northwest of the North Atlantic (18), and the Mediterranean Sea (5), are the bioregions with the highest number of PAs and which also have more than 50% of their area sampled as *Adequate*. While Indo-Pacific seas and Indian Ocean bioregion have over 80% of its protected area without any data among the analyzed databases (i.e. *NR*). (See Appendix D) FAO areas with the largest area categorized as *Adequate* correspond to the northwest Atlantic (20.98%), Northeastern part of the Pacific Ocean (11.78 %), and Western part of the Atlantic Ocean (11.41 %) (Table 3). These FAO areas correspond to regions of the Pacific Ocean (North Pacific, North West Pacific, Mid-tropical N Pacific Ocean and Indo-Pacific seas and Indian Ocean, as well the Gulf of California and Caribbean and Gulf of Mexico). Largest FAO areas with *NR* correspond to the Antarctic part of the Pacific Ocean, the Antarctic part of Atlantic Ocean and Southeastern part of the Atlantic Ocean in the Southern Ocean, Offshore S Atlantic and Southern Africa.

**Table 2:**
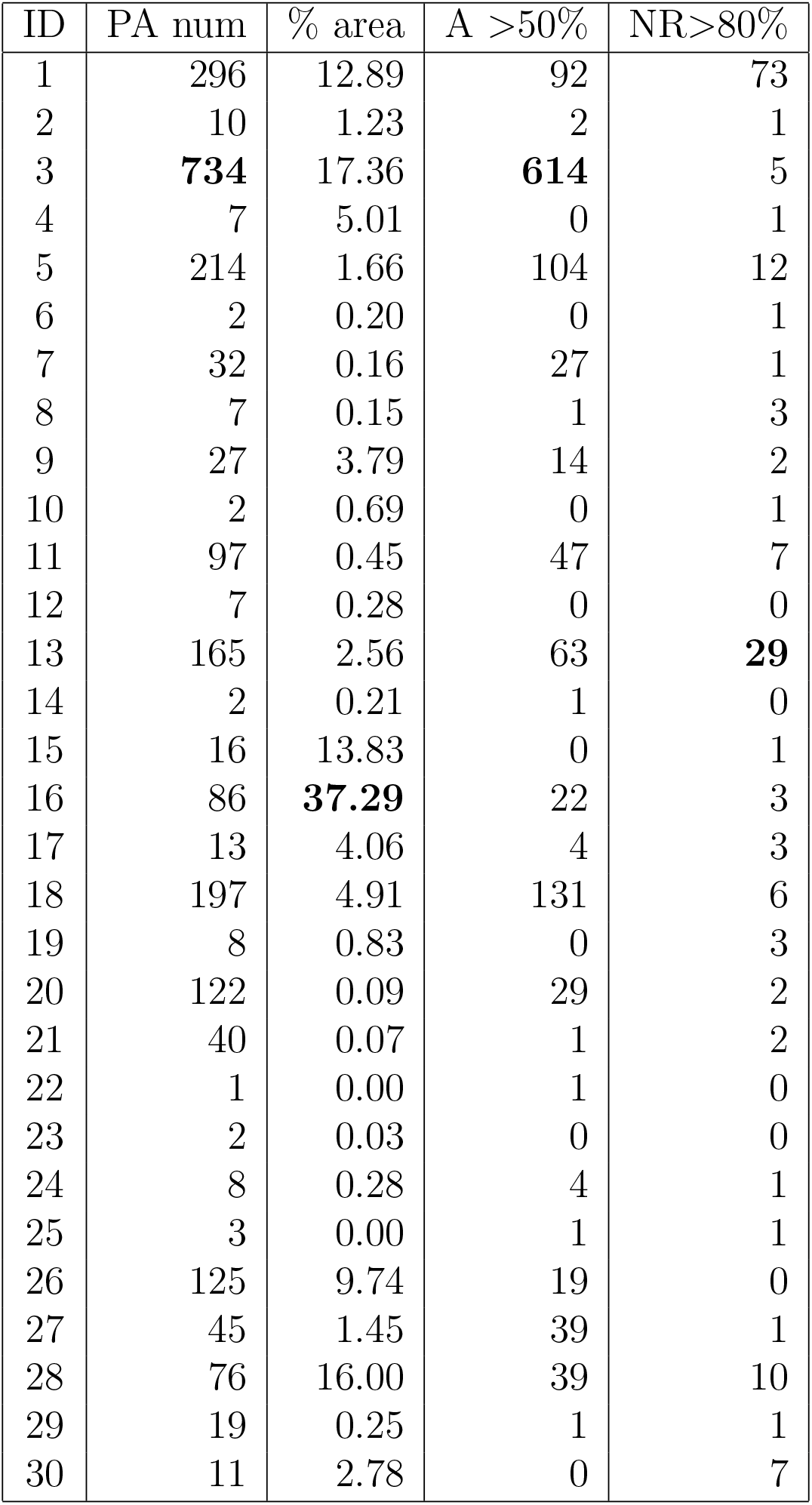
Protected area overlap and SRI. “ID” is the identification number given to each bioregion. “PA num” is the number of protected areas (PA). “P”, the percentage of the surface area covered by PA. “A*>*50%” is the number of protected areas with more than 50% of its surface area classified as *Adequate*. “NR*>* 80%” is the number of PAs with a surface area larger than 80% with *No Records*. The highest values are highlighted in bold face.

| ID | PA num | % area | A >50% | NR>80% |
| --- | --- | --- | --- | --- |
| 1 | 296 | 12.89 | 92 | 73 |
| 2 | 10 | 1.23 | 2 | 1 |
| 3 | **734** | 17.36 | **614** | 5 |
| 4 | 7 | 5.01 | 0 | 1 |
| 5 | 214 | 1.66 | 104 | 12 |
| 6 | 2 | 0.20 | 0 | 1 |
| 7 | 32 | 0.16 | 27 | 1 |
| 8 | 7 | 0.15 | 1 | 3 |
| 9 | 27 | 3.79 | 14 | 2 |
| 10 | 2 | 0.69 | 0 | 1 |
| 11 | 97 | 0.45 | 47 | 7 |
| 12 | 7 | 0.28 | 0 | 0 |
| 13 | 165 | 2.56 | 63 | **29** |
| 14 | 2 | 0.21 | 1 | 0 |
| 15 | 16 | 13.83 | 0 | 1 |
| 16 | 86 | **37.29** | 22 | 3 |
| 17 | 13 | 4.06 | 4 | 3 |
| 18 | 197 | 4.91 | 131 | 6 |
| 19 | 8 | 0.83 | 0 | 3 |
| 20 | 122 | 0.09 | 29 | 2 |
| 21 | 40 | 0.07 | 1 | 2 |
| 22 | 1 | 0.00 | 1 | 0 |
| 23 | 2 | 0.03 | 0 | 0 |
| 24 | 8 | 0.28 | 4 | 1 |
| 25 | 3 | 0.00 | 1 | 1 |
| 26 | 125 | 9.74 | 19 | 0 |
| 27 | 45 | 1.45 | 39 | 1 |
| 28 | 76 | 16.00 | 39 | 10 |
| 29 | 19 | 0.25 | 1 | 1 |
| 30 | 11 | 2.78 | 0 | 7 |

**Table 3:** Results of overlapping FAO fishery exploitation areas and SRI grid. The surface area corresponding to each bioregion, and the percentage of surface area of each classification. Area is in thousands of km2; NR is the percentage of cells with *No Records*; F is the percentage of classified cells with *Few* records; S, the percentage of classified cells with *Sufficient* records, and A, the percentage of classified cells with *Adequate* records. The highest values are highlighted in bold.

| FAO Area Name | Area | NR | F | S | A |
| --- | --- | --- | --- | --- | --- |
| Arctic Ocean | 9.24 | 89.81 | 4.74 | 3.29 | 2.16 |
| Northwestern part of the Atlantic Ocean | 6.28 | 42.48 | 16.91 | 19.62 | **20.98** |
| Northeastern part of the Atlantic Ocean | 14.35 | 62.85 | 13.42 | 14.02 | 9.71 |
| Western part of the Atlantic Ocean | 14.65 | 43.56 | **22.37** | **22.66** | 11.41 |
| Eastern Central part of the Atlantic Ocean | 14.19 | 64.17 | 20.43 | 10.07 | 5.34 |
| Mediterranean Sea and the Black Sea | 2.99 | 57.82 | 19.07 | 12.75 | 10.36 |
| Southwestern part of the Atlantic Ocean | 17.70 | 85.60 | 7.08 | 5.42 | 1.91 |
| Southeastern part of the Atlantic Ocean | 19.07 | 93.34 | 2.13 | 1.61 | 2.93 |
| Antarctic part of the Atlantic Ocean | 12.48 | 95.31 | 2.38 | 1.63 | 0.68 |
| Western part of the Indian Ocean | 29.42 | 79.15 | 10.42 | 4.87 | 5.56 |
| Eastern part of the Indian Ocean | 31.21 | 87.57 | 5.24 | 5.16 | 2.04 |
| Antarctic and Southern of the Indian Ocean | 12.75 | 90.02 | 5.23 | 3.13 | 1.62 |
| Northwestern part of the Pacific Ocean | 21.54 | 80.43 | 10.87 | 6.02 | 2.67 |
| Northeastern part of the Pacific Ocean | 7.63 | 68.73 | 8.53 | 10.96 | 11.78 |
| Northeastern part of the Pacific Ocean | 33.36 | 78.69 | 9.77 | 6.88 | 4.66 |
| Eastern Central part of the Pacific Ocean | **48.25** | 85.09 | 9.01 | 3.47 | 2.43 |
| Southwestern part of the Pacific Ocean | 27.64 | 87.72 | 4.79 | 4.88 | 2.61 |
| Southeastern part of the Pacific Ocean | 30.80 | 92.85 | 4.89 | 1.33 | 0.92 |
| Antarctic part of the Pacific Ocean | 10.23 | **95.61** | 2.28 | 0.71 | 1.40 |

### 3.5. Evaluation of Biases

We evaluated biases for body size, habitat depth, and cultural value for 10,371 marine fish species identified in our database (§3.1).

#### 3.5.1. Body size

The range 0-80 cm is the most frequently occurring size length. Three species stand out with the highest numbers of records, *Scomber scombrus, Lagodon rhomboides* and *Mallotus villosus* with 20,995, 19,563 and 13,609 records respectively. These species are distributed mainly in the Northeast Atlantic (3) and Offshore and Northwest North Atlantic (18) bioregions. While the families that accumulate the greatest number of records correspond to Sparidae, Scombridae and Labridae with 24,837, 21,719 and 21,035 records. These families are mainly distributed in the Caribbean Sea and Gulf of Mexico and the Northeast Atlantic.

#### 3.5.2. Habitat depth

The most frequent depth range is between 0-838 m (i.e. epipelagic and mesopelagic zones), and the species with the highest number of records are *Mola mola, Coryphaena hippurus* and *Lagodon rhomboides* with 21,089, 21,042 and 19,563 occurrences in the databases. These species are distributed mainly around the Caribbean Sea and Gulf of Mexico (11) bioregions, as well as the following bioregions: Offshore and NW North Atlantic (18) and the South Atlantic Coast (21). The families that accumulate a greater number of records correspond to Scombridae, Gadidae, Sparidae with 63,572, 38,876 and 30,041 records. These are mostly distributed in the northern hemisphere. That is, the Caribbean and the Gulf of Mexico (11), Offshore and NW of the North Atlantic (18) and part of the South Atlantic Ocean Coast (21) bioregions.

Figure 5 shows that body size and habitat depth have a negative correlation with the frequency of records.

**Fig. 5:**
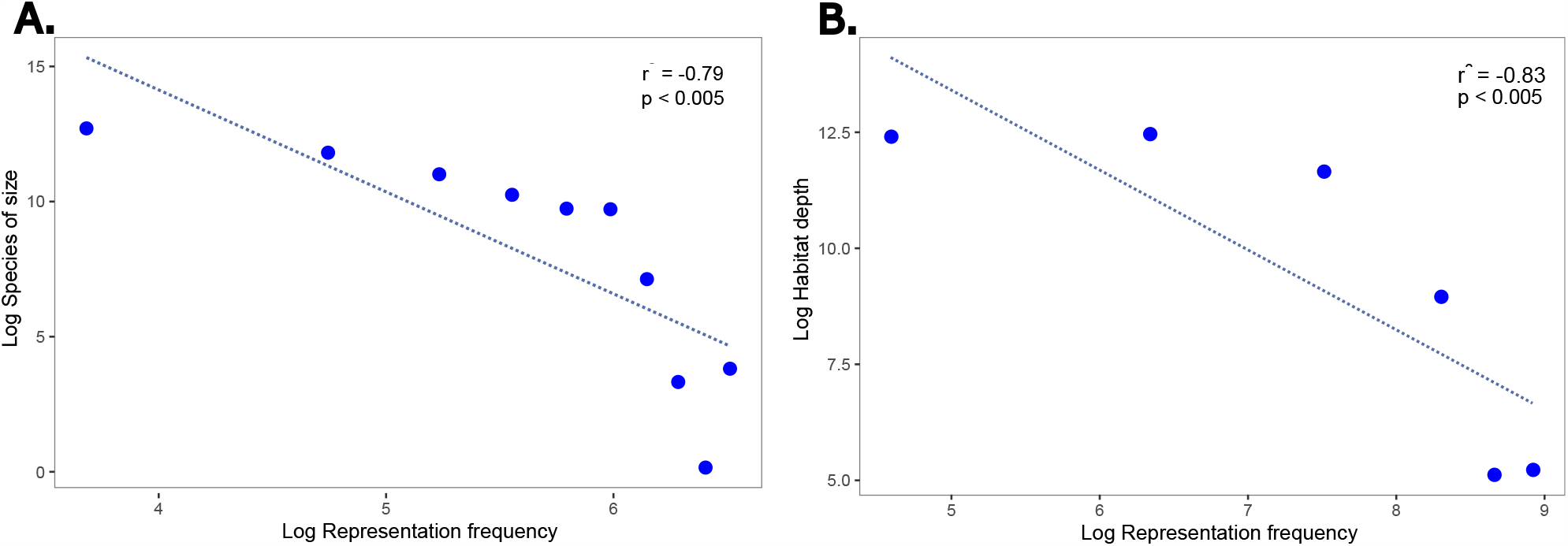
**A**. Relationship between marine fish representation frequency (log_10_ *a*) in GBIF and OBIS and habitat depth. **B**. Relationship between the frequency of representation of marine fishes (log_10_ *a*) in GBIF and OBIS and the size of the species. The dotted line is only there to highlight the negative trend between variables.

#### 3.5.3. Cultural value

Finally, when analyzing the most frequent cultural value represented across our dataset (Fig. 6), “Commercial” use of the species emerges as the most important with a 73.4% among records, followed by the category “No interest” (5.03%), and “Subsistence fishing” (3.08%).

**Fig. 6:**
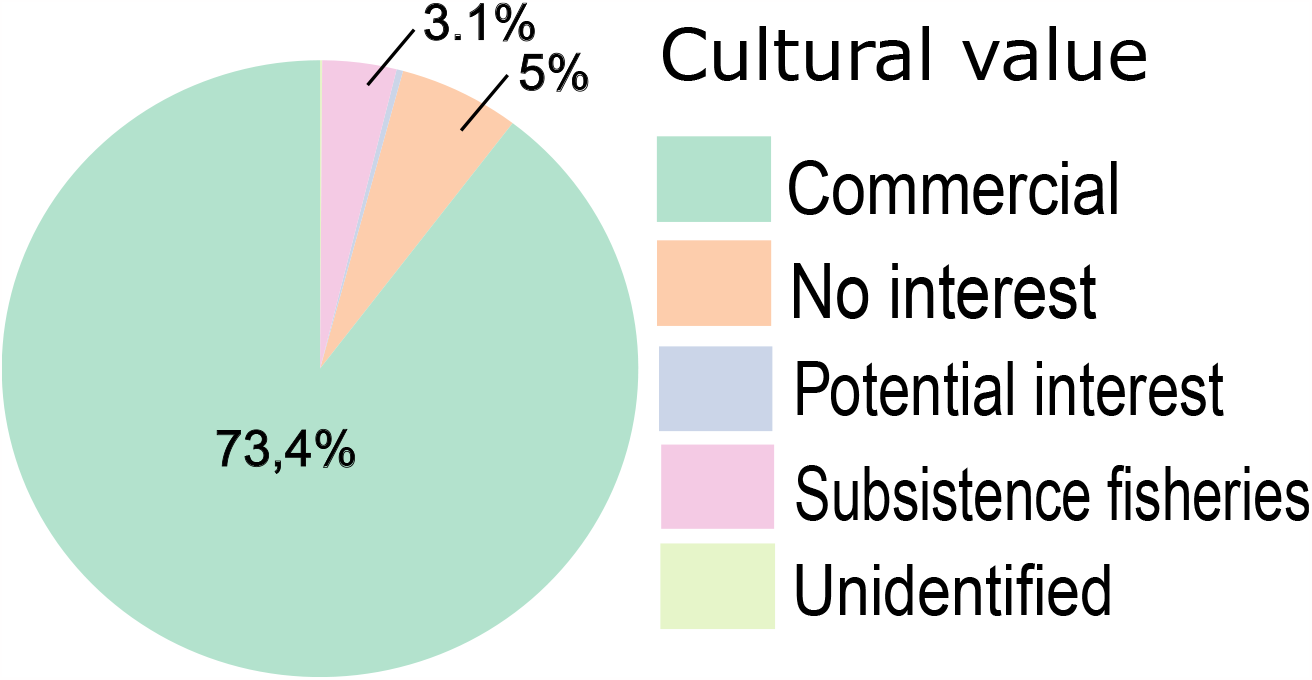
Frequency of marine fish representation in GBIF and OBIS according to importance of cultural use.

## 4. Discussion

Our work provides a methodological framework based on a set of non-parametric estimators to quantify the potential number of species from incidence data (Chao et al., 2009). We used hexagonal grids that fit the geographic reality of marine ecosystems, and we placed special emphasis on cleaning the occurrence data in their taxonomy (Jin and Yang, 2020) and any potential input errors associated with large and massive datasets (Zizka et al., 2020). This led us to focus on only evaluating marine species in the order Actinopterygii (Alò et al., 2021).

Publicly accessible occurrence records are growing rapidly, partly due to the significant advances in ecoinformatics (Lenoir et al., 2020; Oliver et al., 2021). These databases harbor a growing variety of sources, including museum specimens, field observations, acoustic and visual sensors, and citizen science efforts (Amano et al., 2016). However, despite the incredible accumulation of biodiversity records, not all the data is really useful, nor does it represent new insights into the distribution of species (Bayraktarov et al., 2019; Zizka et al., 2020). That is why a systematic evaluation of the integrity and coverage of this information is required (Troia and McManamay, 2017).

There is an extensive bibliography that evaluates the record quality available for different taxonomic groups. Some examples are: legumes on a global scale (Yesson et al., 2007), lepidoptera from Great Britain and woody plants in Panama (Chao et al., 2009), global marine biodiversity (Tittensor et al., 2010), vascular plants in China (Yang et al., 2013), marine fish on a global scale (Mora et al., 2008; García-Roselló et al., 2015), freshwater fish in the USA (Troia and McManamay, 2017; Pelayo-Villamil et al., 2018), and terrestrial mammals on a global scale (Oliver et al., 2021), among many others. Assuming that that not all data available in these repositories are useful for biodiversity analyses, several efforts have proposed parametric and non-parametric estimators for data cleaning and species richness analysis, ModestR (García-Roselló et al., 2013), KnowBr (Lobo et al., 2018), and RWizard (Guisande and Lobo, 2019) among these.

Regarding the units of analysis, here we estimate the species richness at the grid level in order to obtain more uniform results on the distribution of occurrences and avoid overestimating the SRI for marine bioregions (Pelayo-Villamil et al., 2018). In addition, we evaluated two additional grid sizes (5°x5° and 10°x10°), and like other studies, our results show that the coarser the resolution used, the greater the overestimation of the index than richness., and, conversely, the finer the analysis scale, the more deficient the sampling (Tittensor et al., 2010; García-Roselló et al., 2015; Meyer et al., 2015; Troia and McManamay, 2016, 2017). In view of the above, we recommend using grids that really allow observing macro-ecological patterns, especially in coastal regions, which may be underrepresented when using thick and square grids (Pelayo-Villamil et al., 2018).

Considering that more than 40 years of data were analysed, our results demonstrated that on a global scale, the primary marine fish data available on the GBIG and OBIS platforms are still far from being representative and complete. Compared with other studies evaluating the same taxonomic group (Mora et al., 2008; García-Roselló et al., 2015), although we obtained similar macroecological patterns, only 1.14% of the records extracted from both repositories were useful for our analyses. The large percentage of the occurrences presented input errors or did not have the necessary data to generate a reliable analysis (Yesson et al., 2007; García-Roselló et al., 2014).

We also found evidence of strong information biases in the records explored. On the one hand, when analyzing the families and species with the greatest representation, they coincide with groups of fish of commercial interest, demonstrating the existence of **taxonomic bias** of the data (Melo-Merino et al., 2020). This is the case of the families Scombridae, Pleuronectidae and Gadidae, which include species of nutritional importance such as tuna, cod, haddock, among others (Cohen et al., 1990). The same is true for the species with the largest number of records, *H. platessoides* (Pleuronectidae), *C. hippurus* (Coryphaenidae), and *M. mola* (Molidae), the first two are species exploited by the fishing industry, with the exception of sunfish (*M. mola*) which has a wide distribution and is mostly associated with scientific and recreational interest (Pope et al., 2010).

The unequal contribution of data at the spatial level is another factor that must be considered to work with data available on ecoinformatic platforms. There is a clear preference for certain regions and/or ecosystems as a result of **geographical bias**. The literature indicates that the highest data contribution rates correspond to developed countries (Yesson et al., 2007; Chandler et al., 2017), and those coastal regions with better road connectivity (Chandler et al., 2017; Melo-Merino et al., 2020). This information uncertainty is also particularly prevalent in under-sampled marine habitats, such as the deep sea (Webb et al., 2010). Our results coincide with what is described in the literature, regardless of the size of the grid that was used to generate the analysis, the bioregions that include the Atlantic, the Caribbean and the Gulf of Mexico, and the Baltic Sea are the regions with the highest number of area sampled as *Adequate* associated mainly with coastal areas. While the bioregions that include the South and Southeast Pacific (including the southern coast of South America), the Southern Ocean, and the Arctic Seas are the regions with the least spatial representativeness of records, the proportion of cells without records (*NR*) exceeds 90%. This large area without samples will make any attempt to describe species richness and distribution highly unreliable in these bioegions (Yang et al., 2013; Troia and McManamay, 2017). The marine regions that include the water column, the seabed and the subsoil beyond the limits of jurisdiction of countries cover almost half of the Earth’s surface and support a great abundance and diversity of life (Visalli et al., 2020). Even so, when examining the marine ichthyofauna occurrence data, these represent the least sampled areas.

Finally, the **time bias** of the data is also present in our study. Diametric differences in species identification and sampling methodologies over the decades have resulted in the production of databases of variable quality. However, the current era is characterized by more accurate data thanks to improvements in individual capture and identification tools (Costello et al., 2015; Jin and Yang, 2020). For these reasons our approach considers occurrence records since 1980, however, the coverage of occurrence data is uneven over time when comparing between marine bioregions. Despite evaluating more than four decades of data, still 46% of marine bioregions have insufficient sampling efforts. Not surprisingly, the Caribbean and Gulf of Mexico (11) bioregion is the region with the largest input of data, demonstrating once again that geographic sampling bias has strong effects on spatial predictions of species richness (Yang et al., 2013). Future sampling efforts should focus on those bioregions corresponding to low or equatorial latitudes, areas where biogeographic studies show that marine biodiversity is concentrated (Costello and Chaudhary, 2017).

All the biases that we have described, added to the inherent problems in data capture, foster and deepen various information gaps that affect the effective spatio-temporal quantification of biodiversity (Magurran and McGill, 2011). In this study, we have overlapped our estimates of species richness with the global marine protected areas declared up to the beginning of the year 2022 (UNEP-WCMC and IUCN, 2022), and the areas of fishing exploitation reported by the FAO (FAO, 2014). This exercise demonstrates how important it is to have public databases that can faithfully reflect the taxonomic and biogeographical knowledge available for each region of the world (Pelayo-Villamil et al., 2018). Our results indicate that Northeastern Atlantic (3) bioregion has the largest number of protected areas (n=734), of which the vast majority (n=614) have over 50% of their area *Adequate* sampled. However, this bioregion is not considered a marine biodiversity hotspot, compared to the Coral Sea and New Zealand bioregions, which have a larger area protected by marine conservation areas (Ramírez et al., 2017). However, we found a low proportion of well-sampled cells in both regions, demonstrating the existence of important information gaps, at least for fish of the order Actinopterygii. We emphasize the need to correct these information gaps so that conservation efforts that seek the implementation of new marine protection areas can have reliable data so as not to underestimate the biodiversity of species (Sala et al., 2021).

In the same way, by overlapping the bioregions with the fishing exploitation zones, we determined that the North Pacific (7) North West Pacific (29), Mid-tropical N Pacific Ocean (9) and Indo-Pacific seas and Indian Ocean (13) bioregions, as well the Gulf of California (21) and Caribbean and Gulf of Mexico (11) are the regions with the highest representation of the data and where fishing activity is concentrated. According to (Kroodsma et al., 2018), the area corresponding to the central Atlantic and Northeast Pacific present little intense fishing effort, while the regions associated with the Northeast Atlantic, the Northeast Atlantic (Europe) regions, and the Northwest Pacific are known to have a huge fishing development and that is where fishing efforts are concentrated worldwide. The southeastern Atlantic Ocean (FAO area 47 and 88), part of the Pacific Ocean (FAO area 88) and Antarctica (FAO area 48 and 88) are the regions with the highest percentage of cells without records (*NR*= *>*93 %). When compared with the findings of (Kroodsma et al., 2018), these areas agree with the “holes” without fishing effort data, which is explained by the geographical remoteness and the lack of technological development necessary for the fisheries to extend to new domains (Visalli et al., 2020). This limits both the exploitation of marine resources and the collection of data.

The hypotheses that we evaluated in this work were necessary to understand what the data collection trends have been and to be able to take future actions to correct the biases described. Our first hypothesis about the size of the body of the fish was rejected. Small fish species (0-80 cm) are the ones that accumulate the largest number of records, among which *S. scombrus* (Scombridae), *L. rhomboides* (Sparidae), *M. villosus* (Osmeridae), are *<*50 cm species that stand out for presenting the largest number of records, and, in addition, they are distributed in the best sampled regions (Mediterranean Sea, Gulf of Mexico and the Caribbean, and the Atlantic Ocean). The size of the fish is inversely proportional to the abundance and, therefore, to the frequency of human use, both scientific and commercial (Pauly and Palomares, 2005).This difference in the sampling effort generates an evident overrepresentation of the smaller species and therefore deepens the taxonomic bias. The hypothesis about the depth of the habitat is accepted, at less depth there is a greater representation of species of marine fish. The pelagic zone has a high concentration of data and effectively corresponds to shallow regions and therefore easily accessible, which generates all the conditions for data collection (Melo-Merino et al., 2020).It has been pointed out that the concentration of species decreases as the depth of the ocean increases, however, it is precisely these areas that have been least sampled and where there is the greatest probability of discovering new species (Costello and Chaudhary, 2017). This demonstrates the need to concentrate efforts on the deeper regions of the water column (mesopelagic, bathyal, and abyssal) for a more equitable representation of marine ecosystems. Finally, the hypothesis of the use of the species is also accepted. The species of marine fish that have a more beneficial or lucrative use for humans are better represented in the analyzed databases. We believe that this is related to the fact that the fishing industry is one of the main sources of information for platforms such as OBIS (Zhang and Grassle, 2002).

Today, marine ecosystems and their biodiversity face the great challenges of climate change and the impact of human activity, especially those species considered key food resources for survival (Hollowed et al., 2013; Ramírez et al., 2017; O’Hara et al., 2021). It is necessary to focus and strengthen the study of those areas with very few or no records, since the descriptions of the geographic ranges of the species and their temporal dynamics are fundamental measures for the evaluation of the real state of biodiversity (Lenoir et al., 2020; Oliver et al., 2021). Having more reliable data will allow effective conservation actions to be implemented.

## Acnlowledgement

Funding for this research was provided by the National Agency of Research and Development of Chile through project FONDECYT Regular #11211490 to HS. We thank professor Ricardo Giesecke for valuable comments on earlier versions of this manuscript.

## Appendix A. The database

Table A.4 below shows the data loss for each criterion that we have used to clean our database. We downloaded 71,670,596 records from GBIG and OBIS. Only 820,004 records were useful for our analyses.

**Table A.4:** Criteria for filtering occurrence data from GBIF and OBIS using bioregions.

| Database state | Number of records |
| --- | --- |
| Original records from GBIF and OBIS | 71,670,596 |
| Data curation (following Zizka et al. (2020)) | 5,380,439 |
| Taxonomically filtered data | 5,007,322 |
| Deletion of data outside the native range | 820,004 |

Files of the 10,371 marine fish species and their attributes (body size, habitat depth, and cultural value) from FishBase may be found in the GitHub project page of this manuscript: http://github.com/vapizarro/stp_fishes

## Appendix B. Grids resolutions

For spatial representation analysis we evaluated two additional spatial resolutions (5°× 5° = 3,021 cells, and 10°× 10° = 958 cells). Table B.5 contains the results of this analysis for these grids. We have also mapped these results (see Figure B.7), to understand how the effect of spatial resolution on the evaluation of biodiversity macropatterns. Finally, we also plot the frequency of cells for each SRI category for the three grid sizes (R1=1°× 1°; R5=5°× 5°; R10=10°× 10°) to understand how the data is distributed in our analyses (see Figure B.8)

**Fig. B.7:**
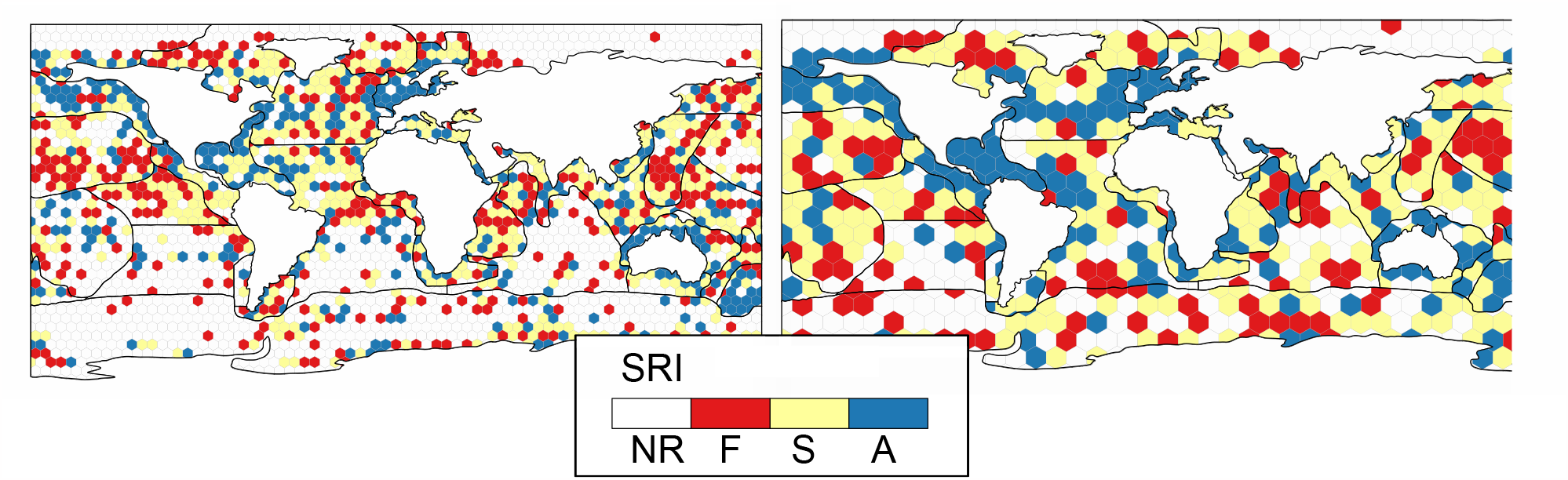
Spatial representativeness index (SRI) mapping of cells of size: A=5°× 5°; B=10°× 10°. The categorization of the cells corresponds to the level reached by the SRI, where SRI ¿ 0.85: Amount of data *Adequate* for the representation of species richness (“A”); SRI=0.60-0.85: Amount of data can be considered *Sufficient* (“S”); SRI=0-0.60: Amount of records *Few* (“F”); and SRI = NA: cells with no records (“NR”).

## Appendix C. Bioregions slopes

We evaluated the slopes of the last 10% of the accumulation curves of each bioregion in our temporal representation analysis. Table C.6 shows the result for each bioregion.

**Fig. B.8:**
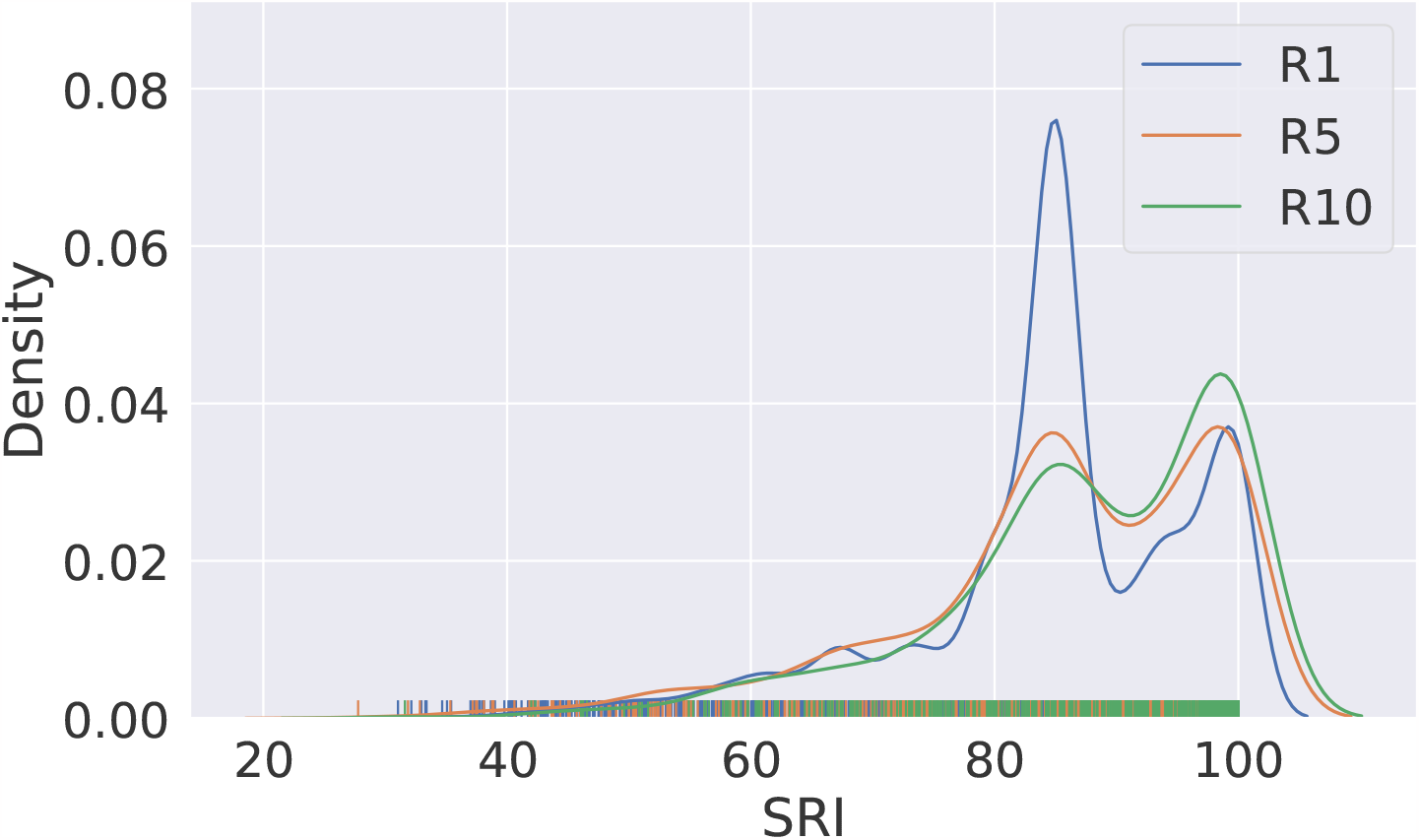
Density probability distribution of SRI in three grids of different sizes: R1 = 1°× 1° (blue line); R5= 5°× 5° (red line); and R10 = 10°× 10° (yellow line).

## Appendix D. Supplementary Material: GAP Analysis

We plotted the percentage of surface with marine protected areas of each bioregion (Fig D.9), and the percentage of cells of each FAO Area for each category of SRI value (Fig D.10).

**Fig. D.9:**
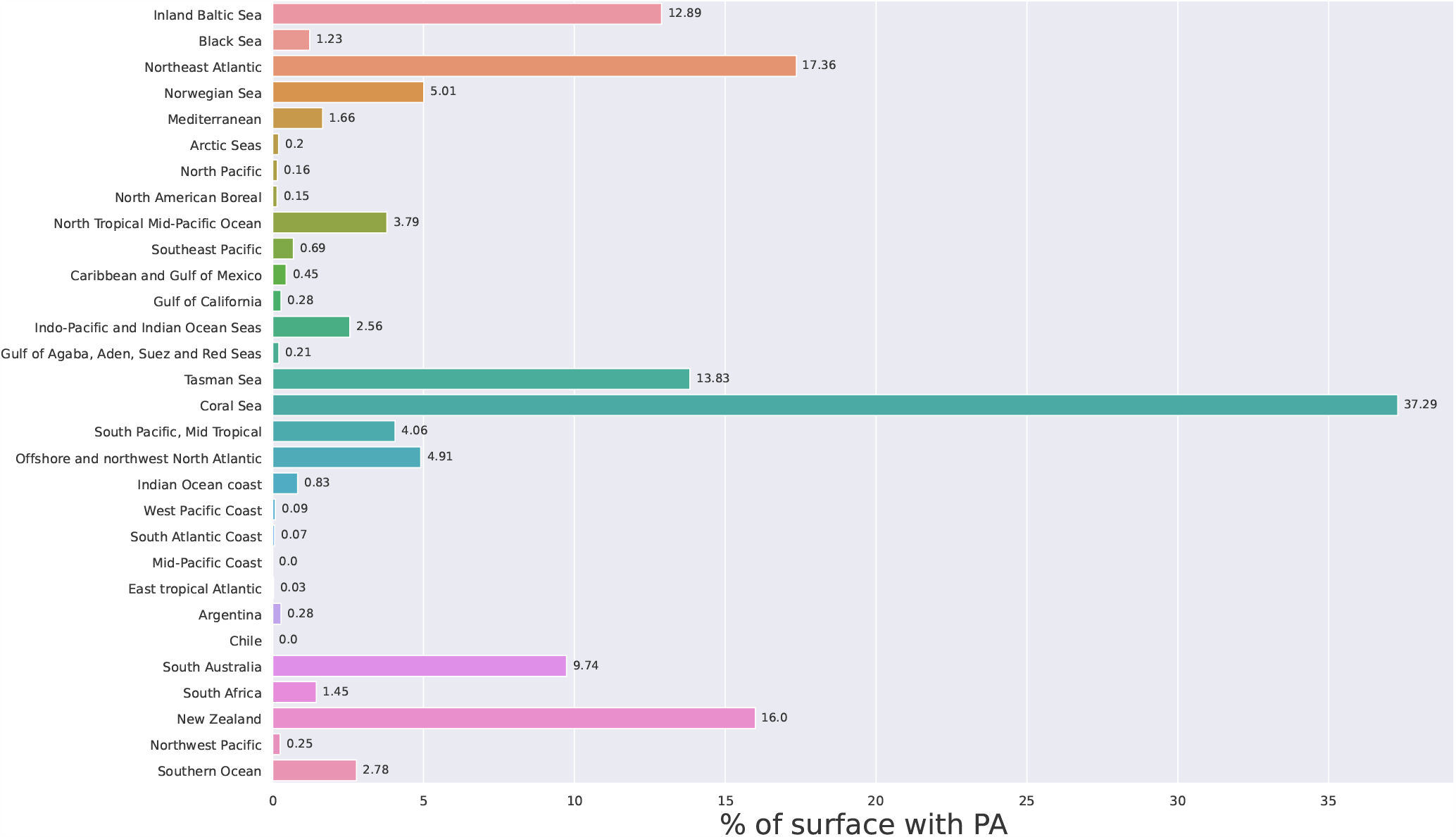
Percentage of surface with marine protected areas by bioregions.

**Fig. D.10:**
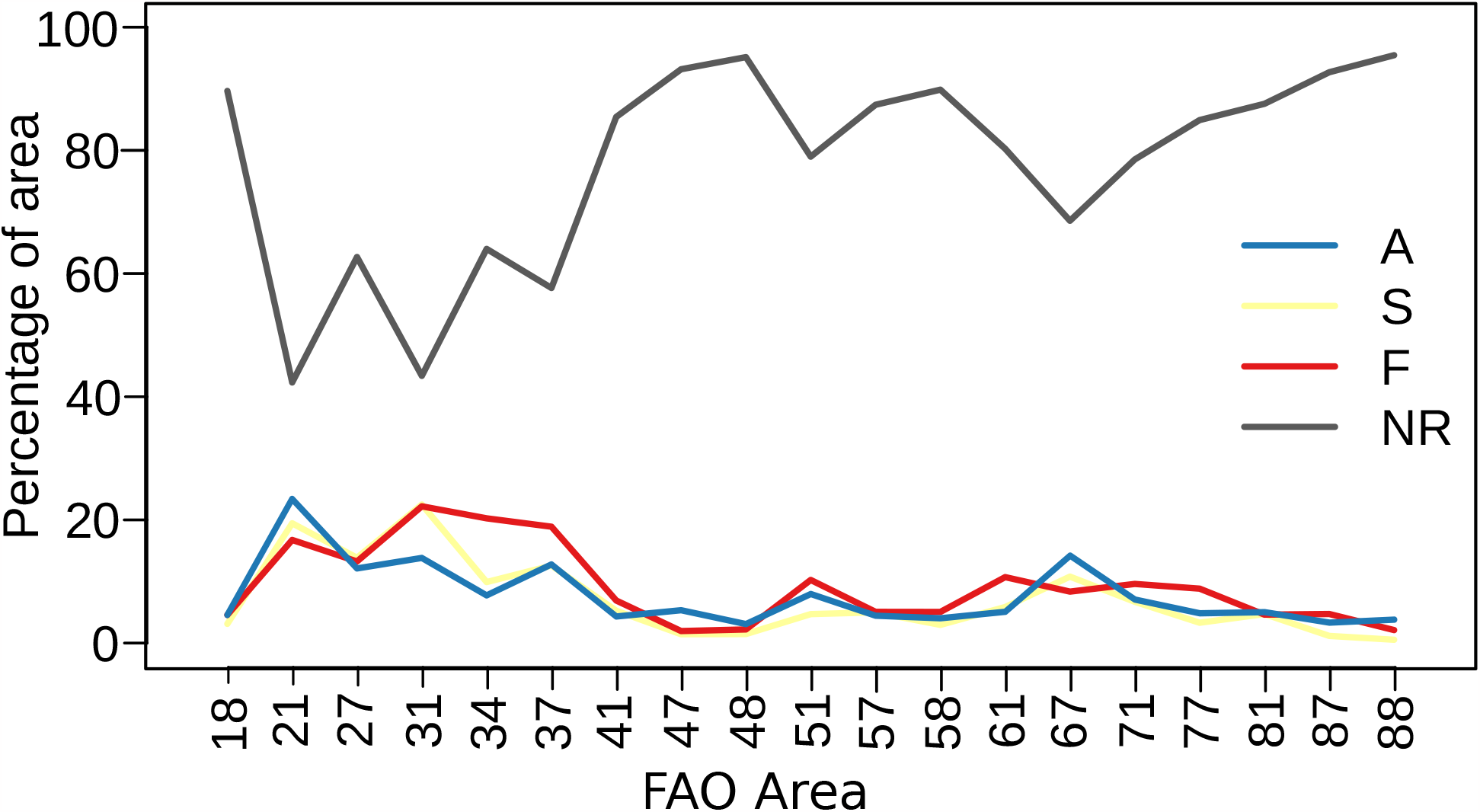
Percentage of cells of each FAO Area for each category of SRI value. Amount of data *Adequate* for the representation of species richness (“A”); SRI=0.60-0.85: Amount of data can be considered *Sufficient* (“S”); SRI=0-0.60: Amount of records *Few* (“F”); and SRI = NA: cells with no records (“NR”).

**Table B.5:** Surface area as a percentage of each bioregion (ID) for every SRI category for each of the three grid sizes (5°×5°; R10=10°× 10°). Amount of data *Adequate* for the representation of species richness (“A”); SRI=0.60-0.85: Amount of data can be considered *Sufficient* (“S”); SRI=0-0.60: Amount of records *Few* (“F”); and SRI = NA: cells with no records (“NR”).

| ID | 5° × 5° |  |  |  | 10° × 10° |  |  |  |
| --- | --- | --- | --- | --- | --- | --- | --- | --- |
|  | NR | F | S | A | NR | F | S | A |
| 1 | 19.38 | 3.52 | 22.18 | 54.92 | 0.17 | 0.00 | 0.00 | 99.83 |
| 2 | 10.22 | 67.79 | 22.00 | 0.00 | 0.24 | 79.80 | 13.41 | 6.55 |
| 3 | 14.17 | 5.86 | 3.68 | 76.30 | 17.26 | 0.00 | 1.65 | 81.09 |
| 4 | 8.11 | 18.15 | 49.78 | 23.96 | 24.40 | 0.00 | 56.89 | 18.71 |
| 5 | 17.61 | 11.41 | 51.06 | 19.91 | 0.76 | 21.01 | 29.87 | 48.37 |
| 6 | 72.22 | 13.59 | 7.01 | 7.18 | 57.95 | 14.53 | 10.67 | 16.85 |
| 7 | 22.68 | 19.18 | 30.95 | 27.26 | 7.23 | 5.57 | 42.67 | 44.52 |
| 8 | 28.29 | 32.88 | 28.34 | 10.49 | 10.75 | 38.44 | 49.14 | 1.68 |
| 9 | 38.10 | 36.07 | 20.03 | 5.80 | 8.97 | 49.76 | 30.88 | 10.39 |
| 10 | 70.33 | 14.72 | 8.21 | 6.73 | 55.39 | 17.91 | 15.12 | 11.58 |
| 11 | 15.36 | 0.00 | 37.46 | 47.18 | 3.84 | 0.00 | 10.04 | 86.11 |
| 12 | 21.88 | 17.33 | 49.92 | 10.87 | 2.44 | 9.04 | 53.22 | 35.30 |
| 13 | 15.01 | 34.79 | 32.35 | 17.85 | 8.84 | 25.58 | 48.64 | 16.94 |
| 14 | 8.31 | 0.00 | 62.35 | 29.35 | 66.52 | 0.00 | 0.00 | 33.48 |
| 15 | 17.06 | 37.36 | 29.94 | 15.64 | 3.42 | 15.88 | 66.66 | 14.04 |
| 16 | 18.45 | 16.45 | 30.57 | 34.53 | 18.82 | 0.00 | 17.13 | 64.06 |
| 17 | 56.51 | 15.47 | 15.27 | 12.75 | 10.25 | 25.29 | 48.27 | 16.19 |
| 18 | 15.10 | 27.12 | 37.68 | 20.17 | 16.20 | 14.33 | 29.37 | 40.10 |
| 19 | 64.31 | 20.32 | 8.07 | 7.30 | 36.75 | 34.23 | 22.00 | 7.01 |
| 20 | 12.21 | 47.20 | 33.32 | 7.27 | 10.74 | 40.12 | 35.84 | 13.30 |
| 21 | 44.88 | 19.20 | 24.68 | 11.23 | 22.43 | 23.33 | 30.03 | 24.21 |
| 22 | 20.76 | 44.23 | 35.01 | 0.00 | 15.95 | 28.49 | 52.89 | 2.67 |
| 23 | 4.42 | 33.22 | 54.38 | 7.97 | 18.89 | 27.78 | 46.90 | 6.43 |
| 24 | 19.18 | 39.23 | 28.34 | 13.25 | 20.90 | 22.41 | 21.87 | 34.82 |
| 25 | 70.95 | 3.80 | 11.03 | 14.22 | 53.42 | 0.00 | 33.72 | 12.86 |
| 26 | 30.23 | 1.16 | 44.14 | 24.46 | 27.42 | 0.00 | 32.14 | 40.44 |
| 27 | 17.67 | 20.53 | 37.77 | 24.02 | 10.57 | 18.38 | 41.56 | 29.48 |
| 28 | 25.40 | 11.73 | 20.42 | 42.45 | 14.84 | 0.00 | 30.51 | 54.65 |
| 29 | 4.47 | 14.81 | 47.69 | 33.03 | 20.81 | 0.00 | 65.06 | 14.13 |
| 30 | 72.92 | 11.70 | 9.40 | 5.98 | 44.37 | 22.69 | 25.57 | 7.37 |

**Table C.6:**
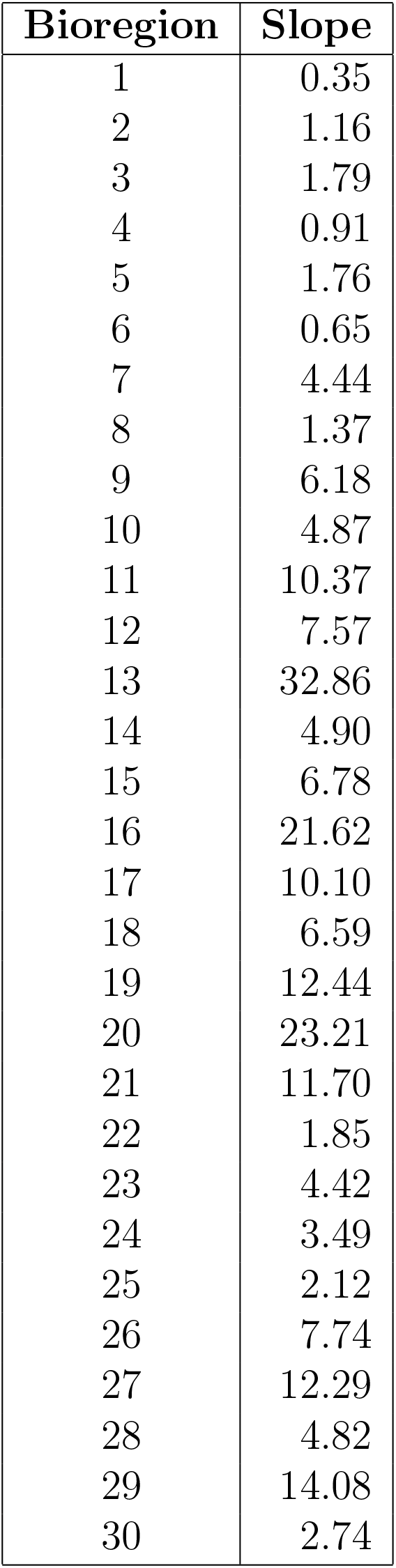
Final slope (10%) of the accumulation curves for each bioregion

